# Coupling of cell growth modulation to asymmetric division and cell cycle regulation in *Caulobacter crescentus*

**DOI:** 10.1101/2024.04.10.587378

**Authors:** Skye Glenn, Wei-Hsiang Lin, Alexandros Papagiannakis, Setsu Kato, Christine Jacobs-Wagner

**Affiliations:** Sarafan Chemistry, Engineering, and Medicine for Human Health Institute, Stanford University, Stanford, CA 94305, USA; Department of Biology, Stanford University, Stanford, CA 94305, USA; Howard Hughes Medical Institute, Stanford University, Stanford, CA 94305, USA; Department of Molecular, Cellular, and Developmental Biology, Yale University, New Haven, CT; Department of Microbiology and Immunology, Stanford School of Medicine, Stanford, CA 94305, USA

**Keywords:** *Caulobacter crescentus*, cell growth, asymmetric division, cell cycle

## Abstract

In proliferating bacteria, growth rate is often assumed to be similar between daughter cells. However, most of our knowledge of cell growth derives from studies on symmetrically dividing bacteria. In many α-proteobacteria, asymmetric division is a normal part of the life cycle, with each division producing daughter cells with different sizes and fates. Here, we demonstrate that the functionally distinct swarmer and stalked daughter cells produced by the model α-proteobacterium *Caulobacter crescentus* have different average growth rates despite sharing an identical genome and environment. The discrepancy in growth rate is due to a growth slowdown during the cell cycle stage preceding DNA replication (the G1 phase), which is predominantly associated with swarmer cell functionality. We also provide evidence that the second messenger (p)ppGpp extends the timing of the G1/swarmer cell stage by slowing growth specifically during the beginning of the cell cycle. Our data further show that cells factor the amount and rate of their growth to control the G1/S transition, allowing them to adjust the time they spend with ecologically important G1-specific traits.

**Significance statement:** Bacterial growth rate modulation is generally associated with changes in genetic make-up or environmental condition. This study demonstrates that the rate of cell growth can also vary between daughter cells and across cell cycle stages under invariant and unstressed environmental conditions. This is illustrated by the asymmetrically dividing α-proteobacterium *Caulobacter crescentus*, which, at each division, produces two functionally distinct daughter cells that differ in average growth rate. This growth rate difference arises from a G1 phase-specific growth slowdown mediated, in part, by the (p)ppGpp alarmone. Altogether, this study showcases the coupling of cell growth modulation to asymmetric division and cell cycle regulation, which may have implications for other α-proteobacteria given their cell cycle similarities with *C. crescentus*.

## Introduction

Cells must grow between divisions to maintain their size. In bacteria, cell growth is typically assumed to be similar between daughter cells and largely constant throughout the cell cycle once normalized for cell size. However, it is unclear whether this assumption applies across bacteria, particularly asymmetrically dividing α-proteobacteria, which tend to exhibit a higher degree of cell cycle control not seen in symmetrically dividing bacteria. Asymmetric divisions and the generation of daughter cells of different size and fate are common, if not the norm, among α-proteobacteria, a large class of diverse organisms that includes free-living and host-associated species (1–7). These bacteria tightly regulate their cell cycles to couple important cellular traits—like dispersal, foraging, adhesion, surface sensing, virulence, or symbiosis—to specific cell cycle phases (8–14).

Asymmetric division and cell cycle controls have been extensively studied in *Caulobacter crescentus*, an oligotrophic α-proteobacterium naturally found in aquatic environments (6). The life cycle of this organism is characterized by a developmental program that is coupled to specific cell cycle events through a sophisticated array of molecular control mechanisms and checkpoints (15, 16). These cell cycle complexities, combined with the absence of overlapping DNA replication cycles, have earned *C. crescentus* a reputation of being more “eukaryote-like”, so much so that the *C. crescentus* cell cycle is typically described with the eukaryotic G1/S/G2 phase convention rather than the common bacterial B/C/D period terminology to refer to the stages before, during, and after chromosome replication (Fig. 1A). At each cell cycle, *C. crescentus* divides asymmetrically to produce two daughter cells of different sizes and functionalities: the smaller “swarmer” cell and the larger “stalked” cell (Fig. 1A) (6, 13). The stalked progeny has a thin polar appendage (stalk) with a sticky holdfast that allows the cell to bind and colonize surfaces (6). This daughter cell is DNA replication-competent at birth, as it initiates a new round of chromosome replication following cell division. Meanwhile, the swarmer daughter cell experiences a G1 phase (B period) in which it is unable to replicate its genome (Fig. 1A), akin to the DNA presynthetic gap of the eukaryotic cell cycle. During this G1 phase, the swarmer cell is endowed with specific cellular functionalities such as motility and surface sensing mediated by the flagellum and pili at a specific pole. Following the G1 phase, the swarmer cell differentiates into a stalked cell through pilus retraction, flagellum ejection, and stalk growth (the swarmer-to-stalked cell transition), and concurrently becomes competent for DNA replication (the G1/S phase transition) (Fig. 1A).

**Figure 1.**
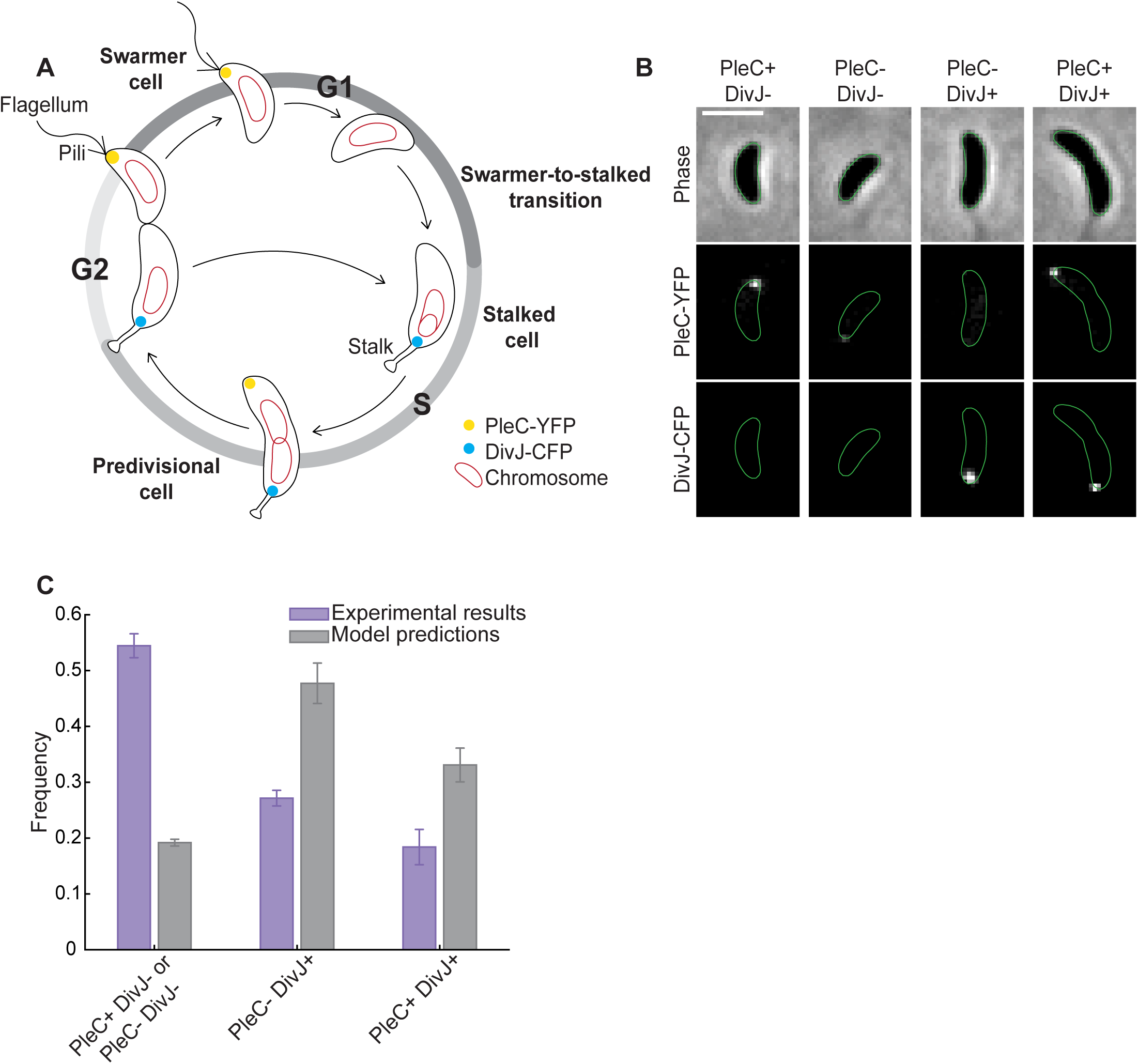
Swarmer cells are overrepresented in populations relative to expectations from a constant growth model. (**A**) Schematic showing a *C. crescentus* life cycle in which the cell cycles of the daughter cells converge. Shown also is the polar localization of the cell cycle markers PleC-YFP and DivJ-CFP. Polar localization of each marker is indicated as PleC+ or DivJ+, whereas the absence of polar localization is labeled as PleC- or DivJ-. (**B**) Representative images of each cell type (strain LS3205) in the Convergent Constant Growth (CCG) model are shown on the right side. Scale bar represents 1 µm. (**C**) Comparison of the expected frequency of each cell type in the population based on the CCG model with experimental data. The model predictions are estimated from the size distributions of each cell type for each replicate. Shown are the mean ± the standard deviation from three independent experiments (n > 1300 cells per experiment).

The cellular and molecular events that control and couple the cell cycle and development in *C. crescentus* have been dissected in significant detail (16–18), which has led to the formulation of several cell cycle models (19–25). However, there remains a lack of clarity around cell growth over the complete life cycle of this organism (or that of any other α-proteobacteria). Technical limitations have largely confined the quantitative study of growth to the stalked progeny (26–32). For this progeny, growth between birth and division is well-described by a single exponential fit (28, 31, 32), indicating that the growth rate is relatively constant between birth and division when normalized by cell size. It should be noted that recent growth analysis of both *C. crescentus* stalked progenies and *Escherichia coli* has brought some nuance to this view, revealing a slight super-exponential growth behavior in both organisms (33). However, there is a lack of quantitative information about the growth of the *C. crescentus* swarmer progeny during its cell cycle. It is possible that the swarmer cell phase corresponds to a growth-arrested stage, with cell growth resuming only after the G1/S phase transition when the swarmer cell differentiates into a stalked cell. Alternatively, swarmer cells may grow at the same rate (normalized by cell size) as their stalked siblings. A state of growth is more consistent with electron micrograph measurements of synchronized cell cultures showing that cell volume increases during the swarmer cell phase (7). However, no growth rate measurement could be extracted by this approach.

Modulation of growth rate during the cell cycle or in response to asymmetric division are known to occur in eukaryotic cells (34–37). Therefore, we set out to determine whether *C. crescentus*, an asymmetrically dividing bacterium with functional traits coupled to cell cycle progression, also modulates cell growth during its life cycle.

## Results

### Characterization of cell type representation in asynchronous populations relative to a simple cell cycle model

To begin examining growth of *C. crescentus* cells in the context of its life cycle, we built a simple model based on ergodic theory (35) to predict the frequency of each cell type in an asynchronous population growing at steady state. In this model (Fig.1A), the predivisional stalked cell (or “predivisional cell”) divides asymmetrically to produce two daughter cells of different sizes. The larger stalked progeny develops into a predivisional cell while the smaller swarmer progeny first proceeds through a developmental phase before converging with its stalked sibling, turning into a predivisional cell as well (Fig. 1A). Our model makes two assumptions: (1) the cell cycle of swarmer and stalked progenies converge to produce predivisional stalked cells and (2) the daughter cells grow at the same average relative rate from birth to division (Fig.1A). Hereafter, we refer to this as the “Convergent Constant Growth” (CCG) model.

Ergodic rate analysis exploits the fact that the fraction of cells at a particular stage is directly related to the time it takes to transit dynamically through that stage. This allows us to predict the expected frequency of each cell type in snapshot images (see Materials and Methods). To identify the cell type inputs for the CCG model, we classified cells (strain LS3205) using two well-described fluorescently-labeled cell cycle markers, DivJ-CFP and PleC-YFP. These fluorescent markers display a reproducible pattern of pole localization at specific cell cycle stages (38), as shown in Fig.1A. Swarmer cells are recognizable by the presence of a fluorescent PleC-YFP focus at a cell pole and the absence of a polar DivJ-CFP fluorescent focus (PleC+ DivJ-) (see example images in Fig. 1B). Additionally, some swarmer cells are expected to have no PleC-YFP or DivJ-CFP focus (PleC-DivJ-) while they are undergoing the swarmer-to-stalked cell transition. Stalked cells are marked by a single DivJ-CFP focus (PleC-DivJ+) whereas predivisional cells generally harbor DivJ-CFP and PleC-YFP foci at opposite poles (PleC+ DivJ+). We took snapshots of the LS3205 strain growing in the standard medium PYE at 30°C and performed cell typing using a spot detection method (see Materials and Methods, Fig. S1A). We found that, under our experimental conditions, swarmer (PleC+ DivJ- and PleC- DivJ-) cells represented 54 ± 2% (mean ± standard deviation or SD, n = three biological replicates) of the cell population, whereas stalked (PleC-DivJ+) and predivisional (PleC+ DivJ+) cells accounted for the remaining 27 ± 1% and 18 ± 3% of the cell population, respectively (Fig.1C). We compared these population statistics to our CCG model predictions based on the cell size distributions for each cell type (Materials and Methods, Fig. S1B). The model, which assumes that growth rate is constant throughout the cell cycle, consistently underestimated the fraction of PleC+ DivJ-/PleC-DivJ-cells at 19 ± 1% while overestimating the fraction of PleC-DivJ+ cells and PleC+ DivJ+ cells at 48 ± 4% and 33 ± 3%, respectively (Chi-square test, p = 0.00 x 10^-100^, Fig. 1C). We envisioned at least two possible reasons for the discrepancies between the model predictions and the experimental results: (i) our methodology, based on fluorescently labeled DivJ or PleC, is not suitable for determining cell type identity and/or (ii) our model assumption of constant growth is incorrect. The latter possibility motivated us to determine the growth rate of both daughter cells from birth to division using timelapse microscopy (Fig. 2A), thereby eliminating reliance on model assumptions.

**Figure 2.**
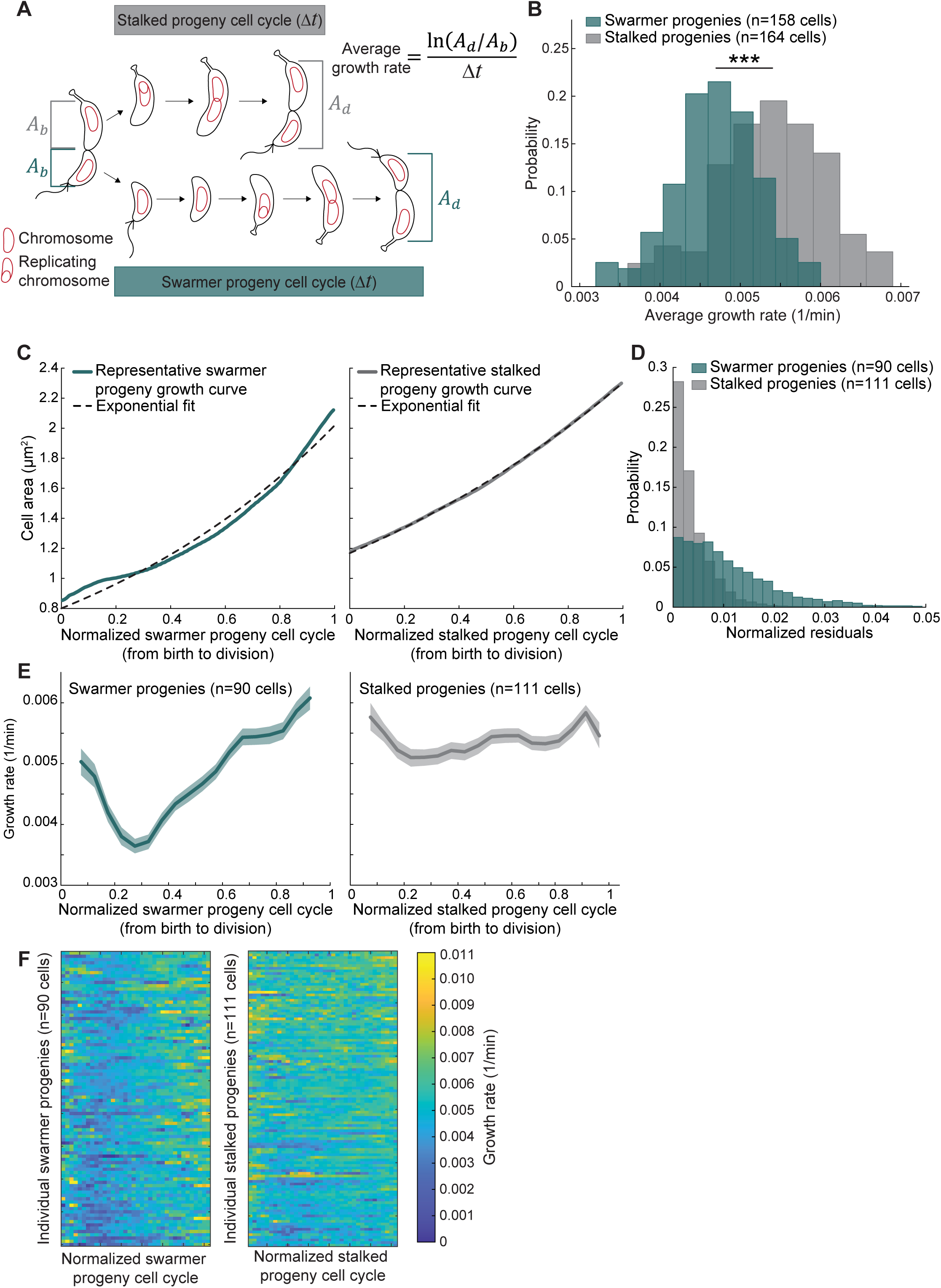
Swarmer and stalked progenies grow at different rates on average. (**A**) Schematic showing a linear representation of the *C. crescentus* life cycle in which the cell cycles of the daughter cells do not converge. Cell area was measured at birth (𝐴𝑏) and division (𝐴𝑑), and average growth rate was calculated over the interdivision time (Δ𝑡). (**B**) Distributions of average growth rate for wildtype (CB15N) swarmer and stalked progenies growing on low-agarose pads. *** indicates p ≤ 0.001 (two-sample Kolmogorov-Smirnov test). (**C**) Representative growth curves of a single swarmer and stalked progeny. The dotted lines are the best exponential fits for each trajectory. (**D**) Plot showing the residuals to exponential fits, normalized for cell area, for all tracked swarmer and stalked progenies. (**E**) Plot showing the growth rate (absolute growth rate normalized for cell area) of all swarmer and stalked progenies, aligned by cell cycle unit. Solid lines and shaded areas denote mean and 95% CI of the mean from bootstrapping, respectively. (**F**) Tempograms of the growth rate for all tracked swarmer and stalked progenies sorted from shortest (top) to longest (bottom) interdivision time.

### Swarmer progenies display slower average growth rate than stalked progenies

Characterizing cell growth over the complete asymmetric life cycle of *C. crescentus* has been hampered by technical limitations. As noted in the introduction, microfluidic devices based on cell immobilization via the holdfast have been effective at tracking the growth of stalked progenies, but not swarmer progenies (26–31). While swarmer cells can be isolated for study using a synchronization technique (39), this method is not well suited for cell growth measurements as the protocol involves growth arrest through cold shock and starvation. Timelapse microscopy on conventional 1% agarose pads containing growth medium allows for visualization of all daughter cells, but results in cell length shortening over generations (40). This is presumably due to confinement by the hard surface and/or local nutrient depletion as this problem is alleviated on a low-agarose (0.3%) pad where *C. crescentus* cells maintain their average size over subsequent divisions (40). Therefore, we performed timelapse phase contrast microscopy on cells growing on low-agarose pads containing PYE medium at 30°C and confirmed that there was no significant difference between cell areas at birth or division between subsequent rounds of division (Fig. S2A). While this method is low throughput, it has the distinct benefit of tracking the cell cycle of both progeny types as the more aqueous environment allows swarmer cells to separate from their mothers at division and swim away from their stalked siblings before occasionally re-attaching to the low-agarose pad nearby (Movies S1 and S2).

From the collected images, we measured the cell areas of swarmer and stalked progenies at birth (𝐴𝑏) and division (𝐴𝑑), defining birth as the frame before the appearance of visible cell separation and division as the frame before the subsequent division (Fig. 2A). We define the swarmer progeny cell cycle as the entirety of the cell cycle between birth and division (from the beginning of the swarmer cell stage to the end of the predivisional cell stage, which includes the swarmer-to-stalked cell transition). From these measurements, we calculated the average growth rate as 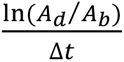 where Δ𝑡 is the interdivision time (Fig. 2A). We found that swarmer progenies grew, on average, about 13% slower (two-sample Kolmogorov-Smirnov test, p = 1.22 x 10^-15^) compared to their stalked siblings (Fig. 2B). Estimating cell volume for cell size measurements yielded similar results as cell area (Fig. S2B). Therefore, we decided to use cell area to calculate growth rate as this metric was directly measured from the two-dimensional microscopy images. The measured difference in average growth rate between daughter cells violates the assumption of constant growth rate in the CCG model, consistent with our population analysis of cell types (Fig. 1C).

### Swarmer progenies modulate growth rate during their cell cycle

To examine growth rate dynamics throughout the swarmer and stalked cell cycles, we extracted the area of 201 cells for all frames (1.5 min imaging interval) between birth and division. As expected (27–29, 31, 32, 40), the increase in cell area of stalked progenies was well-described by a single exponential fit, as illustrated by a representative trace in Fig. 2C (right panel). In contrast, an exponential fit systematically failed to describe the entirety of individual swarmer progeny growth curves, as also exemplified in Fig. 2C (left panel). This deviation resulted in larger fitting residuals for all swarmer progenies compared to stalked progenies (Fig. 2D).

To assess potential systematic changes in growth along single cell cycles, we determined the growth rate of individual cells by calculating the cell area difference between time frames, normalizing for cell area, and plotting it as a function of the normalized cell cycle time from birth (zero) to division (1) (see Materials and Methods). As expected from the good exponential fit to the absolute growth curve (Fig. 2C, right panel), the average growth rate of stalked progenies was largely constant throughout their cell cycle (Fig. 2E, right panel). In striking contrast, the average growth rate of swarmer progenies slowed in the first third of their cell cycle, then increased to eventually reach an average growth rate close to that of the stalked progenies (Fig. 2E, left panel). At the single-cell level, there was considerable heterogeneity (Fig. 2F), as depicted by “tempograms” in which each row represents the growth rate by color scale along a single cell cycle (41). Despite this heterogeneity, the tempogram of swarmer progenies (Fig. 2F, left panel) showed evidence of slower growth rate (shown by a distinctive dark blue trough) in the first half of the cell cycles that was not as apparent in the stalked progenies tempogram (Fig. 2F, right panel). This slower relative growth rate in the first third of the swarmer progeny cell cycle explains the overrepresentation of swarmer cells in asynchronous populations relative to the CCG model predictions (Fig. 1C).

### The growth rate slowdown is associated with the G1 phase

A major cell cycle difference between the two daughter cells of *C. crescentus* regards the G1 phase. The G1 phase is thought to be virtually nonexistent in stalked progenies while swarmer progenies go through an extended G1 phase before they can initiate DNA replication (42) (Fig. 2A). In PYE medium, the swarmer G1 phase has been estimated to last 20-33% of the cell cycle (31, 43, 44), which is within the same range of the observed growth rate decrease in the swarmer cell cycle (Fig. 2E, left panel). As such, we hypothesized that the growth rate change in swarmer progenies is associated with the G1 phase duration. To observe DNA replication in the context of the growth rate change, we performed low-agarose timelapse microscopy on a strain that contains a translational fusion of eYFP to MipZ (Fig. 3A and Movie S3). Fluorescent patches of MipZ-eYFP track the segregation of chromosomal origins of replication (45), which occurs shortly after DNA replication initiation (46). Here, we define the G1 phase as the time between cell birth and the appearance of two MipZ-eYFP patches in the cell (Fig. 3A and Movie S3).

**Figure 3.**
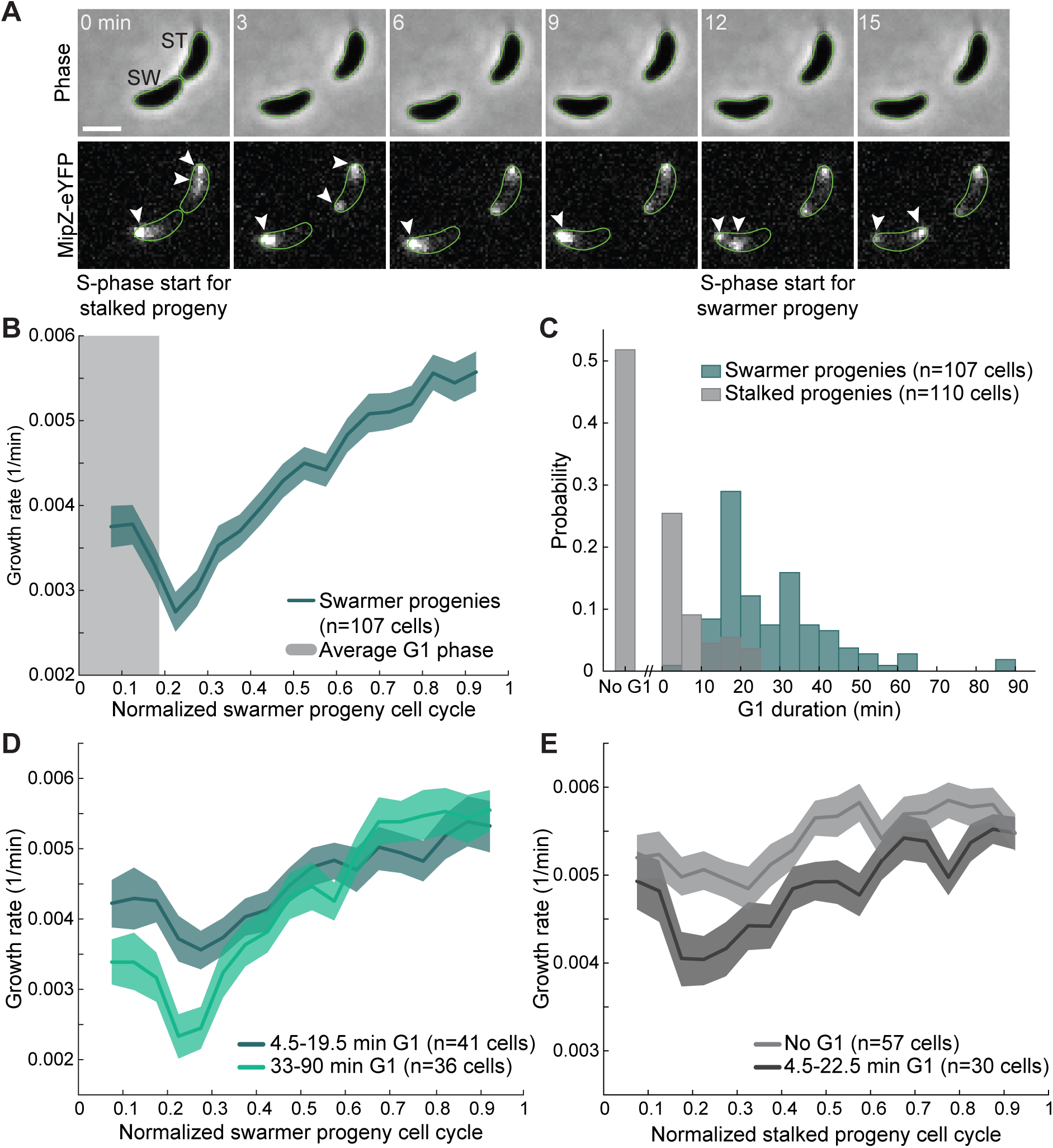
The period of slow growth is associated with time spent in G1 phase. (**A**) Example montage of a stalked (ST) and swarmer (SW) progeny with the replication marker MipZ labeled with eYFP (strain CJW2022). Cell birth (t = 0 min) is defined as the last frame before the daughter cells are clearly separated. White arrows point to fluorescent MipZ-eYFP spots. Scale bar represents 1 µm. (**B**) Plot showing the growth rate of all tracked swarmer progenies expressing a MipZ-eYFP fusion (strain CJW2022). Solid line and shaded area denote mean growth rate and 95% CI of the mean from bootstrapping, respectively. Grey shaded region represents the average G1 phase as measured by MipZ-eYFP segregation. (**C**) Histogram of the absolute duration of the G1 phase across all analyzed swarmer and stalked progenies. (**D**) Plot showing growth rate of swarmer progenies binned according to the indicated time spent in the G1 phase. Solid line and shaded area denote mean and 95% CI of the mean from bootstrapping, respectively. (**E**) Same as panel D but for stalked progenies.

We found that, under our experimental conditions, the average duration of the G1 phase in swarmer progenies approximately corresponded to the first 20% of the cell cycle (∼0.2 cell cycle units), slightly preceding the reversal of growth rate from slowing to accelerating (Fig. 3B). Analysis of individual cells revealed a high degree of variability in G1 duration across swarmer progenies, ranging between 4.5 and 90 min (Fig. 3C). When we binned swarmer progenies by the duration in G1 phase, we found that cells with longer G1 phases have a more pronounced slowing of growth in the early stages of the cell cycle (Fig. 3D).

We also noticed that a portion of the stalked progeny population experienced a short G1 phase (Fig. 3C, Fig. S3A, and Movie S4). By comparing the growth rate of stalked progenies with and without a G1 phase, we found that those with a G1 phase (of at least 4.5 min for comparison with swarmer progenies) also exhibited a slight reduction in growth rate in the early stages of the cell cycle compared to those with no G1 phase (Fig. 3E). This suggests that the slow growth early in the cell cycle is not a phenomenon specific to cells with swarmer identity, but rather a more general feature of experiencing a phase without DNA replication. As such, we refer to the period of slow growth that occurs at the beginning of the cell cycle as G1-associated slow growth, or the “G1 slowdown” for brevity.

### Loss of (p)ppGpp attenuates the G1 slowdown

The second messengers guanosine tetraphosphate and pentaphosphate (collectively referred to as (p)ppGpp) are well-known inhibitors of cell growth in bacteria (47, 48), raising the question of whether (p)ppGpp is involved in the observed G1 slowdown. In addition, overproduction of (p)ppGpp in *C. crescentus*—either artificially or through cell starvation—results in a G1-phase arrest (49–51). While the molecular mechanisms are incompletely understood, G1 arrest occurs when (p)ppGpp accumulation stimulates the proteolysis of the DNA replication inhibitor CtrA and blocks the synthesis of the conserved DNA replication initiator DnaA (49, 50, 52, 53). Bulk culture measurements of the *C. crescentus* Δ*spoT* mutant, which cannot make (p)ppGpp, have shown that this (p)ppGpp-devoid mutant has a slightly shorter doubling time in PYE medium than the wildtype strain (52). The *spoT* deletion has also been shown to result in a lower fraction of swarmer cells in steady state populations, suggesting a shorter swarmer developmental stage (52). A general increase in growth rate distributed evenly across the *C. crescentus* life cycle would not yield this result. Therefore, we examined growth over the complete life cycle of the Δ*spoT* strain on low-agarose pads by timelapse imaging to detect possible cell cycle-specific effects.

We confirmed a faster average growth rate of the Δ*spoT* strain at the single-cell level using low-agarose timelapse imaging (Fig. 4A, two-sample Kolmogorov-Smirnov test, p = 3.61 x 10^-12^). By extracting complete single cell growth trajectories, we found that Δ*spoT* swarmer progenies exhibit an attenuated G1 slowdown compared to the wildtype parent strain (see dashed line, Fig. 4B). This effect appeared confined to early stages of the cell cycle, as Δ*spoT* swarmer progenies did *not* grow faster than wildtype by the end of the cell cycle (see dotted line, Fig. 4B). Furthermore, the growth rate of Δ*spoT* stalked progenies was relatively similar to that of wildtype stalked progenies (Fig. 4C).

**Figure 4.**
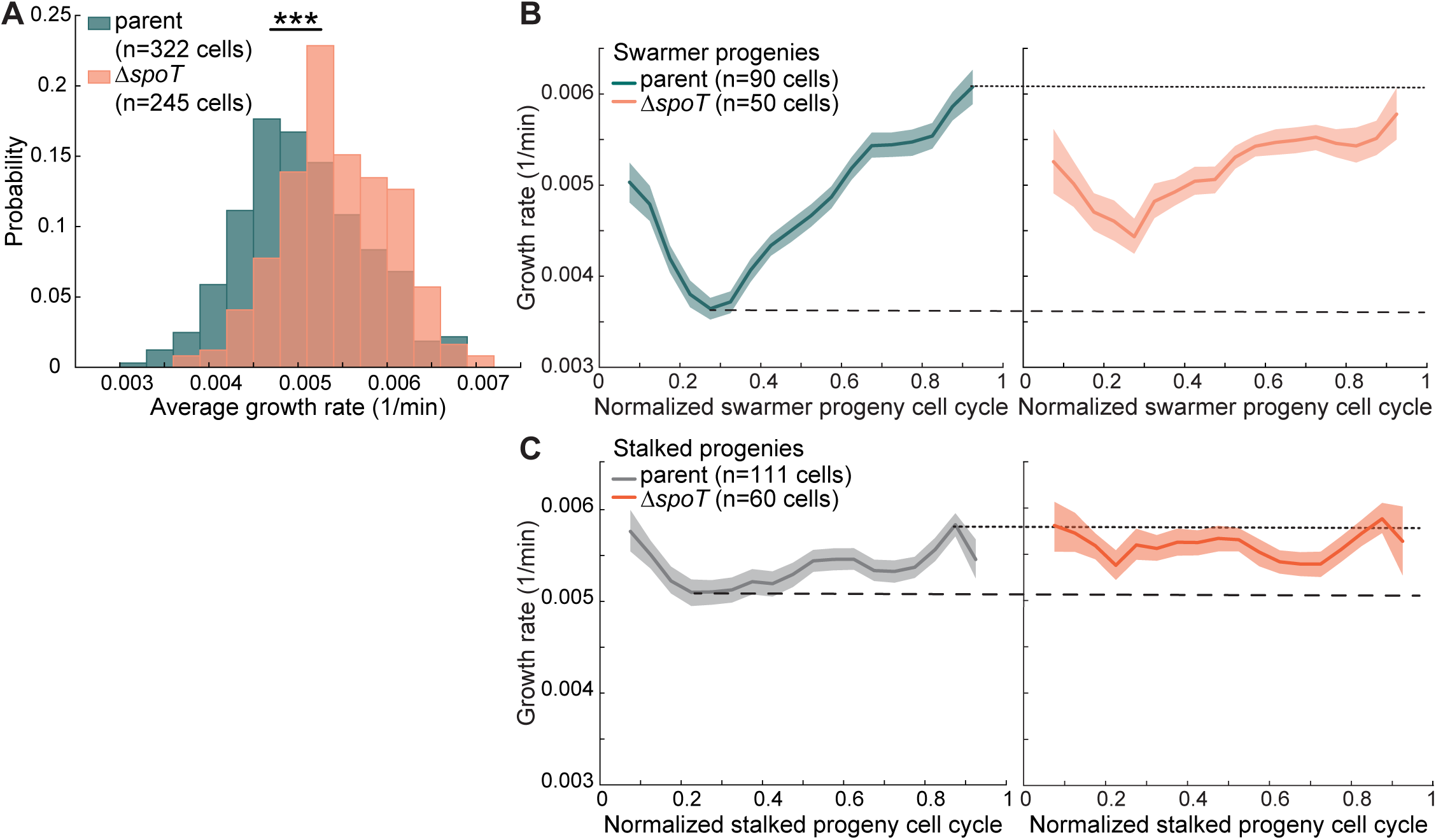
Loss of (p)ppGpp increases the population growth rate by attenuating the G1 slowdown. (**A**) Distributions of average growth rates calculated between cell birth and division of swarmer and stalked progenies (combined) from the wildtype (parent) strain (CB15N) and the Δ*spoT* strain (CJW7364). *** indicates p ≤ 0.001 (two-sample Kolmogorov-Smirnov test). Data for the wildtype strain are the same as those shown in Fig. 2B and are replicated only for visual comparison. (**B**) Plots comparing growth rate between swarmer progenies of the parent (CB15N) and Δ*spoT* (CJW7364) strains as a function of the cell cycle. The data for the parent strain are the same as those shown in Fig. 2E and are replicated only for visual comparison. Solid lines and shaded areas denote mean and 95% CI of the mean from bootstrapping, respectively. Dotted and dashed lines are guides showing the upper and lower limits of mean growth rate for the parent (CB15N) strain, respectively. (**C**) Same as panel B but for stalked progenies.

These results support the notion that (p)ppGpp contributes, at least in part, to the G1 slowdown observed in wildtype swarmer progenies. This also potentially explains the lower proportion of swarmer cells observed in Δ*spoT* population measurements (52). If cells transit faster through the G1 phase while growing at a similar rate at other points in the life cycle, they might undergo the G1/S transitions sooner. As a result, swarmer cells would be less represented in the population. This raised the question of whether cell growth regulates the G1/S transition.

### The G1/S transition is linked to cell growth

The G1/S transition in *C. crescentus* has been extensively studied at the molecular level. This has led to the identification of key molecular players and an essential cascade of biochemical events that culminate in the initiation of DNA replication (henceforth referred to as the G1/S regulatory cascade) (17, 54). The association between the growth slowdown after birth and the duration of the G1 phase (Fig. 3C-E) suggested a role for cell growth or size in the G1/S transition, which, to our knowledge, has not been considered in cell cycle modeling studies (19–25).

At the phenomenological level, the initiation of DNA replication (and the coordinated swarmer-to-stalked cell transition) may happen at a constant time after cell birth (timer), at a critical cell size (sizer), or after a constant cell size has been added (adder) (55). A timer would be expected if the regulation of the G1/S transition depended only on the time it took for molecular events (such as the G1/S regulatory cascade) to occur following cell birth. In such a case, the G1 phase duration should be independent of cell size at birth. However, this is inconsistent with the observed dependence of G1 duration on cell area at birth (Fig. 5A). Our MipZ-based method slightly overestimates the timing of DNA replication initiation since the method reports a subsequent step (i.e., the visible separation of the duplicated origins of replication). Nonetheless, such a systematic delay is unlikely to artificially create a cell size dependence.

**Figure 5.**
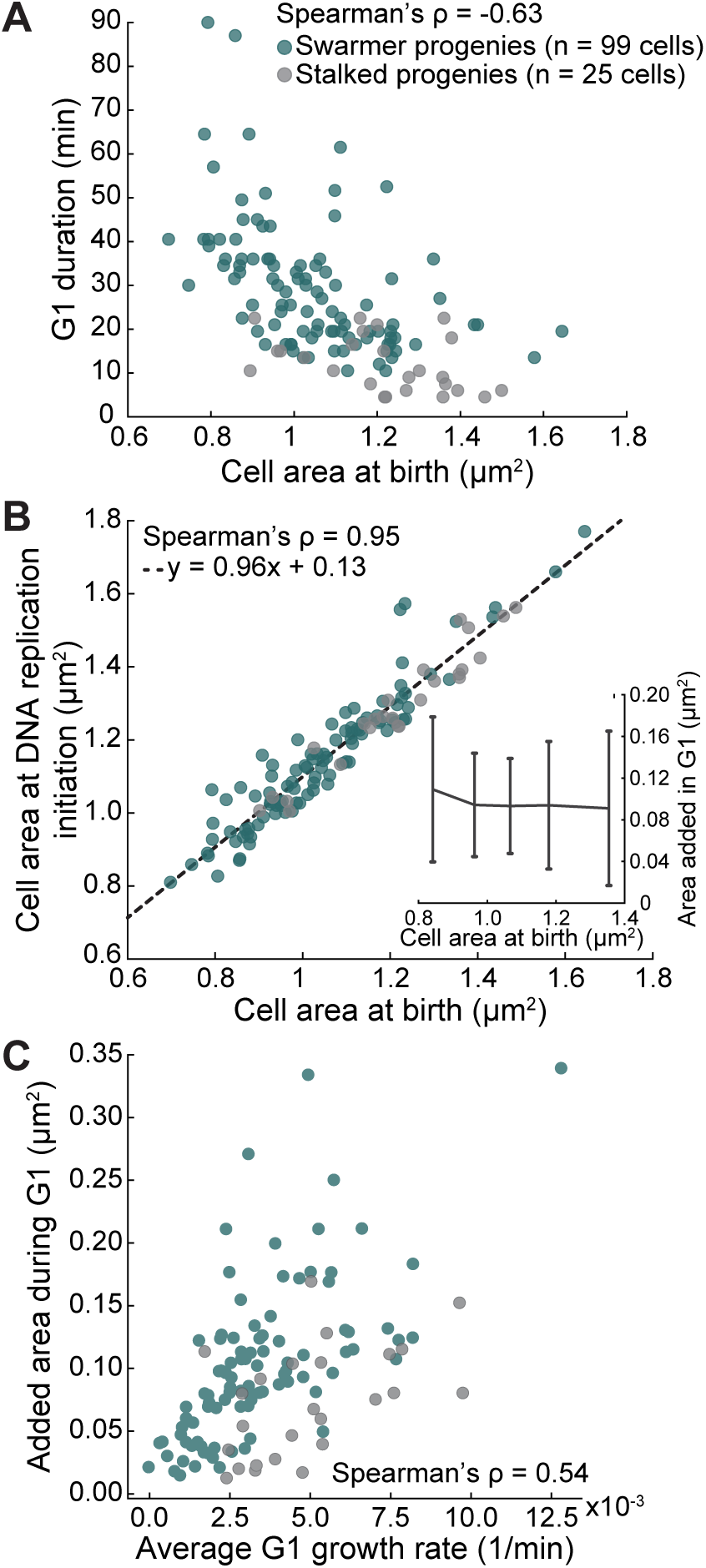
Cell growth contributes to the control of the G1/S transition. (**A**) Scatter plot showing the relationship between cell area at birth and G1 phase duration for the indicated cells (CJW2022) with G1 phase duration ≥ 4.5 min. A minority of cells (9%) with negative values for area increase between birth and DNA replication initiation were excluded. (**B**) Plot showing cell area at birth versus cell area at DNA replication initiation for swarmer and stalked progenies (CJW2022) shown in panel (A). Inset, distributions of added cell area between birth and MipZ-eYFP spot duplication for different bins of cell areas at birth. (**C**) Correlation between the average growth rate during the G1 phase and the added cell area during the same period for the cell populations shown in panel (A).

To distinguish between a G1 sizer and adder, we examined the relationship between cell size at two events, cell birth and DNA replication initiation. A perfect sizer implies that the initiation of DNA replication occurs at a specific cell size regardless of size at birth, yielding a slope of zero. On the other hand, a perfect adder, in which all cells add the same size increment between cell birth and DNA replication initiation regardless of size at birth, would give rise to a slope of one. We observed a strong correlation (Spearman’s correlation = 0.95) and an almost direct proportionality (linear regression slope = 0.96) between cell area at birth and cell area at DNA replication initiation for swarmer and stalked progenies with a G1 phase. This is inconsistent with a sizer and in agreement with an adder whereby cells grow, on average, a similar amount (i.e., add a similar average cell area) before initiating DNA replication, irrespective of their area at birth (inset, Fig. 5B). However, this adder was “noisy” based on the large standard deviation of added cell area across cells (inset, Fig. 5B), with a coefficient of variation (CV) of 0.62 for the combined stalked and swarmer cell populations. This variability in added cell area across cells could be partially explained by differences in average growth rate during the G1 phase, as suggested by the positive correlation (Spearman’s correlation = 0.54) between the two variables (Fig. 5C). That is, fast growing cells (i.e., the cells with a less pronounced G1 slowdown) accumulated more biomass from birth to the onset of DNA replication than slower growing cells. This supports the notion that *C. crescentus* integrates cell growth information to control the G1/S transition.

## Discussion

While *C. crescentus* has been extensively studied with respect to asymmetric cell division, cell development, and intracellular organization for several decades (15–17, 54, 56), a characterization of cell growth over the complete cell cycle of both *C. crescentus* progenies has been missing. Here, we demonstrate that this bacterium modulates its growth in a manner coordinated with the G1 phase. Cell cycle-coordinated growth rate regulation is common in eukaryotes, though the pattern over the cell cycle can vary across cell types (34–36). From a broad perspective, the growth rate pattern of *C. crescentus* swarmer progenies is reminiscent of that of some mammalian cells. Specifically, HeLa cells and retinal pigment epithelial cells exhibit a transient G1 slowdown in growth followed by an increase in growth rate at the G1/S transition (35). *C. crescentus* exhibits checkpoints and cell cycle controls typically not present in symmetrically dividing bacteria (11, 12, 15, 57, 58). As such, *C. crescentus* has long been considered to exhibit more eukaryote-like behavior. Its cell cycle modulation of growth rate described here extends this metaphor into the realm of cell growth.

It is often assumed that the *C. crescentus* daughter cells differ by the presence or absence of a G1 phase. While this accurately reflects the average behavior of swarmer and stalked progenies, this notion is more nuanced at the single-cell level, as stalked progenies can display a short G1 phase (Fig. 3C). It should also be noted that the G1 phase is, by convention, defined to start at cell birth, i.e., after the late predivisional cell separates into two distinct cells. However, our low-agarose timelapse technique, which clearly demarks the time of cell separation with the swarmer progeny movement (Movies S1 and S2), shows that some stalked progenies initiate DNA replication in the late predivisional cell before cell separation (e.g., Fig. S3A). Thus, DNA replication can initiate in the preceding division cycle. This is likely because the timing of membrane invagination is uncoupled during cell division in *C. crescentus*, with the invaginating inner membrane fusing before the outer membrane (59). This uncoupling between inner and outer membranes results in the generation of late predivisional cells with two functionally separated cytoplasmic compartments (11, 60). Therefore, the true start of G1 phase may correspond to the end of cytokinesis (i.e., when one cytoplasm is divided into two in the late predivisional cells) rather than cell separation in *C. crescentus*.

Notwithstanding these details, it is clear that most swarmer progenies display considerably longer G1 phases than stalked progenies (Fig. 3C). As a result, swarmer progenies are, on average, associated with a stronger G1 slowdown during the cell cycle (Fig. 2E), resulting in slower average growth rates relative to their stalked siblings (Fig. 2B). A difference in growth rate between daughter cells may not be unique to *C. crescentus*, as asymmetric divisions appear common among other α-proteobacteria despite their diversity in lifestyle and ecology (3–5, 9, 14, 61). Furthermore, the smaller daughter cells of the plant symbiont *Sinorhizobium meliloti,* the plant pathogen *Agrobacterium tumefaciens,* and the free-living species *Hyphomonas neptunium* initiate DNA replication later relative to their bigger siblings (62–64), similar to what is observed in *C. crescentus*.

The fact that the G1 slowdown can be partially overcome by a Δ*spoT* deletion (Fig. 4) suggests that *C. crescentus* does not use its maximal growth potential during the G1 phase, raising the question of what these cells are prioritizing. The G1 phase has been shown to be essential to the lifestyles of α-proteobacteria as a time in which cells acquire important traits. In *C. crescentus*, the G1 phase of the swarmer cell cycle is associated with motility, chemotaxis, and pili-based surface sensing (65–68). In other α-proteobacteria, the G1 phase is associated with symbiosis, virulence, or dispersal (8, 9, 14, 69). For example, the animal and human pathogen *Brucella abortus* infects host cells primarily during the G1 phase (8). Given the functional importance of this phase, the ability to modulate the time spent in G1 would seem an important factor in the cell’s ability to respond to a changing environment. Indeed, *C. crescentus* swarmer progenies are known to spend more time in the G1 phase upon nutrient limitation or depletion (43, 50–52, 70, 71). Swarmer cells have also been reported to reduce the time of transition to stalked cell identity when the G1-specific pili mechanically sense a surface for attachment (72). G1 duration based on a timer would not offer such flexibility, dooming cells to differentiate after a given amount of time has lapsed regardless of the environmental suitability. On the other hand, a coupling between the G1 phase and cell growth (Fig. 5) would allow cells to integrate information about the growth conditions (i.e., the environment) before committing to the costly process of DNA replication.

How do cells connect nutritional sensing to G1-dependent cellular functionality and growth rate regulation at the molecular level? Our findings, together with past observations, point to the secondary messenger (p)ppGpp as a conserved link. It has been shown that elevated (p)ppGpp levels, generated either artificially through the synthesis of a constitutively active (p)ppGpp synthase or by inducing starvation, lead to an overrepresentation of the swarmer cell type in these populations by extending the G1 phase (49, 50, 52, 70). In *Brucella* spp., where the G1-phase cell is the primary infectious form (8), synthesis of (p)ppGpp is required for proliferation in phagocytic cells and for survival in mice (73, 74). Conversely, overproduction of (p)ppGpp results in an accumulation of G1-phase cells in the population (75). In *Rhizobia* spp., mutants lacking (p)ppGpp are severely impacted in their ability to enter into a symbiotic relationship with the leguminous host plant (76, 77) whereas (p)ppGpp overproduction due to nitrogen deprivation delays the G1/S phase transition (78). At least in *C. crescentus*, these (p)ppGpp-dependent effects are thought to occur by inversely modulating the protein stability of DnaA and CtrA (49, 50, 52, 53), which have opposing effects on DNA replication (79, 80). Our data suggest that (p)ppGpp also affects cell cycle progression through growth rate modulation (Fig. 4B). Similar to wildtype, the Δ*spoT* mutant adds about the same average amount of area (i.e., growth) before transitioning to the S phase under the same growth conditions (Fig. S4A). This mutant achieves this increment of growth sooner because it grows faster during that time (Fig. 4B), resulting in shorter G1 durations (Fig. S4B). Taken together, we propose that (p)ppGpp integrates relevant environmental and intracellular information to influence growth rate, which, in turn, affects the time cells spend in the functionally relevant G1 phase. Other factors must be involved as the G1 slowdown was not completely abrogated in the Δ*spoT* mutant (Fig. 4B).

How is the G1/S phase transition regulated? Significant progress has been made over the years to identify the molecular determinants underlying the G1/S phase and swarmer-to-stalked cell transitions in *C. crescentus*. These two transitions are controlled and coordinated via a cascade of biochemical events and a complex web of temporally and spatially regulated factors that include two-component signal transduction proteins, transcriptional regulators, proteases, scaffolding proteins, polarity factors, and second messengers (16, 17, 81, 82). Growth modulation (Figs. 5 and S4) suggests an additional layer of regulation by which cells control the G1/S phase transition. We hope that our findings will motivate future modeling and experimental studies that will incorporate cell growth modulation as another regulatory dimension to the fascinating life cycle of *C. crescentus*. By extension, it may ultimately become relevant to our understanding of pathogenesis, symbiosis, or other important processes associated with the G1 phase of α-proteobacteria.

### Materials and Methods Bacterial strains and growth

Bacterial strains and their constructions are listed in Table 1. Phage transductions with ɸCR30 and conjugations with S17-1 strains were performed as described previously (83). The plasmid pNPTS138 (M. R. Alley, unpublished) was used for two-step gene replacement with counterselection, as described previously (84). Gibson assemblies were performed using the Gibson Assembly^®^ Master Mix from NEB (85). Primers are listed in Table 2.

**Table 1.** *C. crescentus* strains used in this study.

| Identifier | Genotype | Source |
| --- | --- | --- |
| CJW1472 | CB15N *xylX*::pXGFPC1pXyl-eGFP | Plasmid pXGFPC-1 (89) was electroporated into CJW27. The presence of GFP signal was verified by microscopy after induction with xylose. |
| CJW2022 | CB15N *mipZ*::*mipZ*-eYFP | (87) |
| CB15N | parent strain | (39) |
| CJW3365 | CB15N *rplA(L1)*::pL1-GFPC-1 | (90) |
| CJW7064 | CB15N *spmX*::*spmX*-*mCherry* *rplA(L1)*::pL1-GFPC-1 | φCR30 transducing phage lysate was prepared from CJW3365 and transduced into MT237. The presence of RplA-GFP signal was verified by microscopy. |
| CJW7081 | CB15N *spmX*::*spmX*-*mCherry* pXyl::pXGFPC1pXyl-eGFP | φCR30 transducing phage lysate was prepared from CJW1472 and transduced into MT237. The presence of GFP signal was verified by microscopy after induction with 0.3% xylose. |
| CJW7364 | CB15N Δ*spoT* | pNPTS138 was amplified using primers JSG_104 and JSG_105. Approximately 1 kb of sequence immediately up- and down-stream of *spoT* was amplified from CB15N using primers JSG_106 with JSG_107, and JSG_108 with JSG_109, respectively. The plasmid was assembled by Gibson assembly and electroporated into *E. coli* S17-1 cells. The S17-1 strain was conjugated with CB15N, followed by selection on kanamycin and nalidixic acid. A second cross-over was conducted using the *sacB* counterselectable marker to remove the plasmid sequence. *spoT* deletion was confirmed by PCR amplification using the primers JSG_112 and JSG_113. |
| CJW7370 | CB15N Δ*spoT* *spmX*::*spmX*-*mCherry* | Plasmid described for CJW7364 was conjugated into MT237, followed by selection on kanamycin and nalidixic acid. A second cross-over was |
|  |  | conducted using the *sacB* counterselectable marker to remove the plasmid sequence. *spoT* deletion was confirmed by PCR amplification using the primers JSG_112 and JSG_113. |
| CJW7376 | CB15N $\Delta spoT$ *spmX::spmX-mCherry* *rplA(L1)::pL1-GFPC-1* | ϕCR30 transducing phage lysate was prepared from CJW3365 and transduced into CJW7370. The presence of RplA-GFP signal was verified by microscopy. |
| CJW7751 | CB15N $\Delta spoT$ *mipZ::mipZ-eYFP* | pNPTS138 was amplified using primers JSG_104 and JSG_105. Approximately 1 kb of sequence immediately up- and down-stream of *spoT* was amplified from CB15N using primers JSG_106 with JSG_107, and JSG_108 with JSG_109, respectively. The plasmid was assembled by Gibson assembly and electroporated into *E. coli* S17-1 cells. The S17-1 strain was conjugated with CJW2022, followed by selection on kanamycin and nalidixic acid. A second cross-over was conducted using the *sacB* counterselectable marker to remove the plasmid sequence. *spoT* deletion was confirmed by PCR amplification using the primers JSG_112 and JSG_113. |
| LS3205 | CB15N *divJ-CFP* *pleC-YFP* | (38) |
| MT237 | CB15N *spmX::spmX-mCherry* | (91) |

**Table 2.**
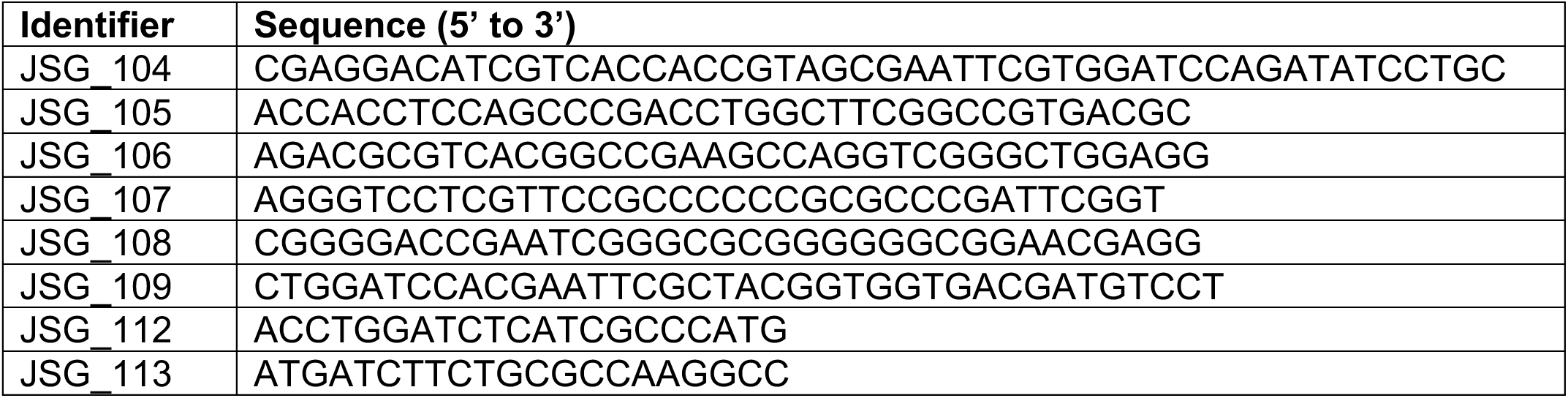
Primers used in this study.

All strains were grown at 30°C in PYE (2 g/L bacto-peptone, 1 g/L yeast extract, 1 mM MgSO4, 0.5 mM CaCl2). To achieve steady-state growth, cells were grown in overnight (∼18 h) cultures then diluted at least 1:10,000 in fresh PYE and grown to exponential phase (OD_660_ < 0.3) in an incubator shaker (∼ 300 rpm, New Brunswick Innova 44) prior to microscopy.

Growth curves for the parent (CB15N) and Δ*spoT* (CJW7364) strains were performed in a 96-well plate using a Synergy2 microplate reader (BioTek). Stationary phase precultures were subcultured into fresh PYE medium to a dilution of 1:10,000 before loading onto the plate. Cultures were grown at 30°C for 33 h with shaking, taking OD_660_ measurements every 2 min.

### Epifluorescence microscopy

Phase contrast and epifluorescence images were acquired on a Nikon Ti-E microscope equipped with a Perfect Focus System, a 100x Plan Apo λ 1.45 NA oil immersion objective, a motorized stage, and an Orca-Flash 4.0 V2 142 CMOS camera (Hamamatsu). For timelapse imaging, an objective heating ring was used set to 30°C and phase images were collected every 1.5 min. When appropriate, YFP images were collected every 3 min. Chroma filter sets were used to acquire fluorescence images: YFP (excitation ET500/20x, dichroic T515lp, emission ET535/30m), CFP (excitation ET436/20x, dichroic T455lp, emission ET480/40 m), GFP (excitation ET470/40x, dichroic T495lpxr, emission ET525/50m) and mCherry (excitation ET560/40x, dichroic T585lp, emission ET630/75m). The microscope was controlled using NIS-Elements AR. When performing timelapse imaging, cells were imaged on 0.3% agarose pads made with PYE. For snapshot imaging, 1% agarose pads made with PYE were used.

### Image processing and analysis

Cell meshes were derived from phase contrast using the open-source image analysis software Oufti (86). Cell size characteristics (e.g., area and volume) were taken directly from Oufti or approximated using the MATLAB function Extract_Extended_Cell_Properties.m (56). Growth rate and associated cell characteristics were calculated using the MATLAB script cell_trajectory_analysis.m. In brief, cell area growth curves were smoothed over a 12-frame sliding-average window (equivalent to 18 min) and the difference in the cell area between consecutive frames was calculated. The relative growth rate was calculated by dividing the absolute growth rate by the cell area from the first of the consecutive frames. Residuals were calculated as the difference between the cell area trajectory and a first-degree polynomial fit to the log-transformed cell area trajectory for each cell. To approximate the timing of DNA replication initiation from timelapse images, the first frame at which the MipZ-eYFP cloud appeared expanded or separated into two distinct clouds (45, 87) was manually assigned as the end of the G1 phase. G1-specific cell characteristics (e.g., G1 duration) were calculated using the MATLAB script cell_trajectory_analysis.m.

### Quantification and statistical analysis

All sample statistics and correlation coefficients were calculated using built-in MATLAB or Excel functions. For the Kolmogorov-Smirnov test, p-values were calculated using the built-in MATLAB function kstest2. For the Chi-square test, the p-value was calculated using the built-in Excel function CHISQ-TEST, with the “actual range” being the frequencies of measured cell types and the “expected range” being the frequencies of cell types calculated by the CCG model (see next section) assuming that all cell types have the same growth rate. Spearman’s rank correlation coefficient was used to assess the degree of association between variables. The built-in MATLAB function fitlm was used to estimate the standard error of the y-intercept of the linear regression to the relationship between cell area at birth and cell area at DNA replication initiation.

### Convergent Constant Growth (CCG) model

*Mathematical model on stage-age distribution and growth rate:* Assuming convergence of the swarmer and stalked cell cycles, the *C. crescentus* cell cycle can be partitioned into three stages (Fig. 1A): swarmer cells, stalked cells, and predivisional cells. Each stage was identified by a combination of the presence or absence of cell cycle markers labeled with fluorescent proteins, PleC-YFP and DivJ-CFP, as indicated in Fig. 1A. In this section, we derive the mathematical formulae of the CCG model using the proposed three-stage *C. crescentus* life cycle. We start with the ideal condition that swarmer, stalked, and predivisional stalked cell cycle stages have constant time intervals 𝜏_1_, 𝜏_2_, and 𝜏_3_. Once this time interval is reached, the cell enters the next cell cycle stage. We define *stage-age* as the time after the cell enters a new cell cycle stage. In this notation, for swarmer, stalked, and predivisional stages, the cell stage-ages range between [0, 𝜏_1_], [0, 𝜏_2_], and [0, 𝜏_3_], respectively.

For the swarmer, stalked, and predivisional cell cycle stages, we define 𝑔_1_(𝑠), 𝑔_2_(𝑠), and 𝑔_3_(𝑠) to be the probability distributions of finding a cell at stage-age 𝑠. Since the probability of all stages sums to one, we have

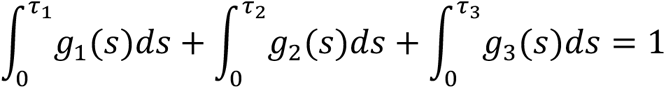

Now, supposing the entire population of cells has a bulk exponential growth rate 𝜆, we use a known result (see Scherbaum and Rasch, 1957, also explained below) that all stage-age distributions 𝑔_*k*_(𝑠) follow an exponential decay with rate 𝜆; that is,

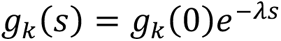

Based on the *C. crescentus* cell cycle model that assumes constant growth and cell size convergence between progenies, we have the following boundary conditions between stage-age distributions:

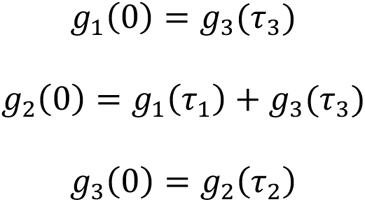

Using the formulae above with some algebraic calculation, we arrive at the relation 𝑒^-𝜆(𝜏_2_+𝜏_3_)^ (1+*e*^-𝜆𝜏_1_^) = 1 and the normalization of age distribution gives 𝑔_1_(0) = 𝜆, 𝑔_2_(0) = 𝜆𝑒^𝜆(𝜏_2_+𝜏_3_)^, 𝑔_3_(0) = 𝜆𝑒^𝜏_3_^. Hence, the fraction of cells at stage *k*, defined by 𝑀_*k*_, can be calculated from the integration 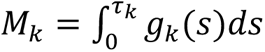. As a result, we have

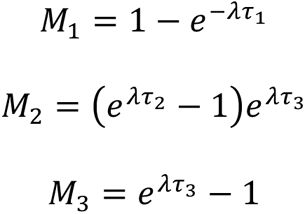

This gives the relation between *M_k_*. and 𝜏*_k_*. Now, we can calculate the stage-specific growth rate, denoted as 𝜇*_k_*. for *k* = 1, 2, 3. If we use cell area as the reference of cell growth, the definition of stage-specific growth rate follows the relation

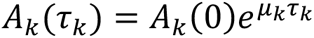

where 𝐴*_k_*(𝑠) is the cell area of cell cycle stage *k* with stage-age *s*. Using the above formula, we have

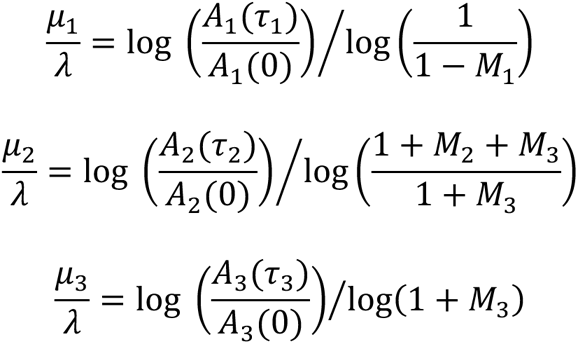

For the CCG model where we assume that all cell stages have the same growth rate, this requires 𝜇_1_ = 𝜇_2_ = 𝜇_3_ = 𝜆. Substituting this equality condition into the above equation, we have

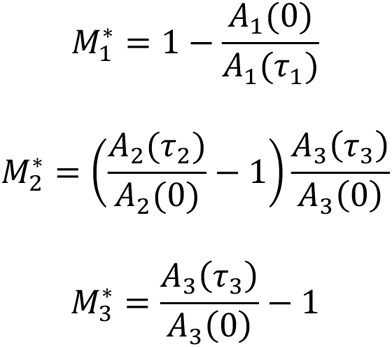

where 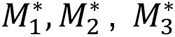 are the expected fractions of cells in the swarmer, stalked, and predivisional stages, respectively.

### Cell typing by image analysis

Cells carrying PleC-YFP and DivJ-CFP (strain LS3205) were spotted on 1% agarose pads and imaged by phase contrast and fluorescence microscopy. A typical image dataset included 1000-2000 cells. Three biological replicates were collected on different days from different cultures. A customized MATLAB code was used for analysis as described below:

(a) Cell detection: Cell detection was based on a segmentation algorithm using phase contrast images. Briefly, the phase contrast images were inverted and subjected to a customized adaptive thresholding algorithm (Mask_adap_thr.m). The algorithm parameter was chosen such that the contour of segmentation matches the cell boundaries in the image based on visual inspection. Cell objects that were located at image boundaries or were too small to represent a cell (area < 0.55 µm^2^) were removed from subsequent analysis. During cell recognition, we simultaneously assigned a cell identifier and measured the cell area.

(b) Cell midline: An additional customized algorithm (cell_coordinate.m) was then used for detecting the cell midline. The cell midline serves as a curvilinear coordinate used to specify the cell region where the fluorescence markers are located.

(c) Background removal: Fluorescence images have non-uniform fluorescence background, which is higher at the center of the images and lower at the boundaries. To remove this non-uniform background, a binary mask was generated from the phase contrast channel to specify the “cell-free region” on each image. For each fluorescence channel, fluorescence data of the cell-free region was fitted to a quadratic function

𝐵𝑔(𝑥, 𝑦) = 𝑎𝑥^2^ + 𝑏𝑥𝑦 + 𝑐𝑦^2^ + 𝑑𝑥 + 𝑒𝑦 + 𝑓. The fluorescence signal was then obtained using the following formula:

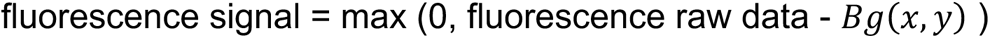

and was used for subsequent analysis.

(d) Fluorescence spot detection: A customized adaptive threshold algorithm (Mask_adap_thr.m) was used to detect the fluorescence spots of PleC-YFP and DivJ-CFP markers. For each channel, the detected spots were subjected to additional filtering criteria: (i) The spot needed to be physically contained within a cell or within two pixels of a cell. (ii) The size and fluorescence intensity of the spot needed to pass a certain threshold. Specifically, all spots were required to have a size greater than three pixels and intensities higher than 200 arbitrary units for PleC-YFP and 300 arbitrary units for DivJ-CFP. (iii) The spot needed to be located at the polar region of the cell. The cell midline was used as a reference coordinate to exclude spots that were located within a 40% fraction of the cell area that corresponds to the mid-section. Spots that satisfied the above conditions were recorded for the following cell typing procedure.

(e) Cell typing: Our image analysis pipeline (double_markers) generated a list of cells with the following associated attributes: (i) cell area, (ii) presence/absence of a PleC-YFP spot, (iii) presence/absence of a DivJ-CFP spot.

(f) Data visualization: A code (sub_save_tile_image_M2) for data visualization of the image analysis pipeline (see Fig. S1A for an example) was developed. Accurate cell segmentation and correct cell typing was verified by manual inspection.

### Parameter extraction

(a) 𝐴*_k_* (0) and 𝐴*_k_*(𝜏*_k_*): In the mathematical model, we assumed no cell-to-cell variability on cell area at each stage. In reality, the ensemble of cells exhibits variability, and the age distributions of cell areas (denoted as 𝑔*_k_*(𝑠) above) overlap, as shown in Fig. S1B. To estimate single values from distributions, we normalized distributions of 𝑔_1_(𝑠), 𝑔_2_(𝑠), 𝑔_3_(𝑠), and 𝑔_1_(𝜃_sw_𝑠), as shown in Fig. S1B. Here, 𝜃_67_ ≡ 0.45 is the division ratio between newborn swarmer cells and predivisional cells, which is similar to previously reported values (13, 32, 40). The cell size distribution of 𝑔_1_(𝑠) was smaller than 𝑔_2_(𝑠), consistent with the asymmetric cell division of *C. crescentus*.

We used the position where these curves intersect to estimate the average starting and ending cell areas. The rationale is as follows: if the cell-to-cell variability is zero, these curves will not intersect, instead forming cell size distributions with boundaries at 𝐴_1_(0), 𝐴_2_(0), 𝐴_3_(0), and 𝐴_1_(0) + 𝐴_2_(0) = 𝐴_3_(𝜏_3_). When these cell size distributions are convolved with noise, they become mutually overlapping functions where the intersections correspond to the original boundary positions. Namely, we use the intersecting positions 𝐴_1_, 𝐴_2_, 𝐴_3_(see Fig. S1B for notation) to estimate the original boundary positions:

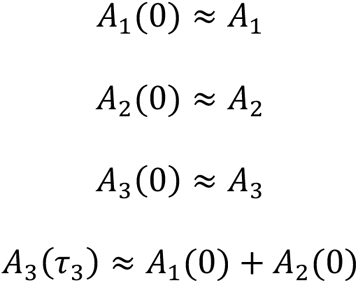

(b) 𝑀*_k_*: The relative frequencies of cells at each stage were obtained directly from cell-typing statistics (see above).

### Mathematical derivation for exponential decay of age distribution

In this section, we derive the equation 𝑔*_k_*.(𝑠) = 𝑔*_k_*.(0)𝑒^-^*^λs^*. For an exponentially growing population, we used 𝑛*_k_*.(𝑡, 𝑠) to denote the number of cells at cell cycle stage 𝑘 at time 𝑡 with cell stage-age 𝑠. The dynamics of 𝑛*_k_*.(𝑡, 𝑠) follow a transport type partial differential equation. Specifically, 𝛿𝑡 denotes a small time interval, and we assume 𝑠 ∈ (0, *𝜏_k_*.) where *𝜏*. is the constant time interval for cell cycle stage 𝑘. Next, we considered a cohort of cells with cell stage-age 𝑠 at time 𝑡. As the time increases from 𝑡 to 𝑡 + 𝛿𝑡, the stage-ages of these cells increase from 𝑠 to 𝑠 + 𝛿𝑡. We assumed that cells do not switch stage (by assuming 𝑠 < *𝜏_k_*.) and that there is no cell death in this model. As such, the number of cells must be the same between 𝑡 and 𝑡 + 𝛿𝑡. Therefore, we have

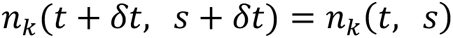

In large populations, we can view 𝑛*_k_*.(𝑡, 𝑠) as a continuous variable of time 𝑡 and cell stage-age 𝑠. If we let 𝛿𝑡 → 0, we have the following partial differential equation

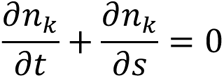

Assuming that the population converges to a stationary distribution 𝑔*_k_*(𝑠) and has bulk growth rate 𝜆, we have 𝑛*_k_*(𝑡, 𝑠) → 𝑔*_k_*(𝑠)𝑒^𝜆*t*^. Substituting this stationary solution into the above partial differential equation, we obtain the relation 𝑔*_k_*(𝑠) = 𝑔*_k_*(0)𝑒^-𝜆*s*^.

## Data and code availability

The data generated by this study are available from the corresponding author (C.J.-W.) upon reasonable request. All original code developed as part of this study has been deposited in the publicly accessible Github code repository (https://github.com/JacobsWagnerLab/published/tree/master/Glenn_et_al_2024).

## Acknowledgements

We would like to thank the Jacobs-Wagner Laboratory for support, discussion, and critical reading of the manuscript. We would also like to thank Dr. Lucy Shapiro and the Shapiro Laboratory group members for sharing strains, as well as for helpful discussions and feedback. C.J.-W. is an investigator of the Howard Hughes Medical Institute.

## Supplementary Figure Legends

**Fig. S1.**
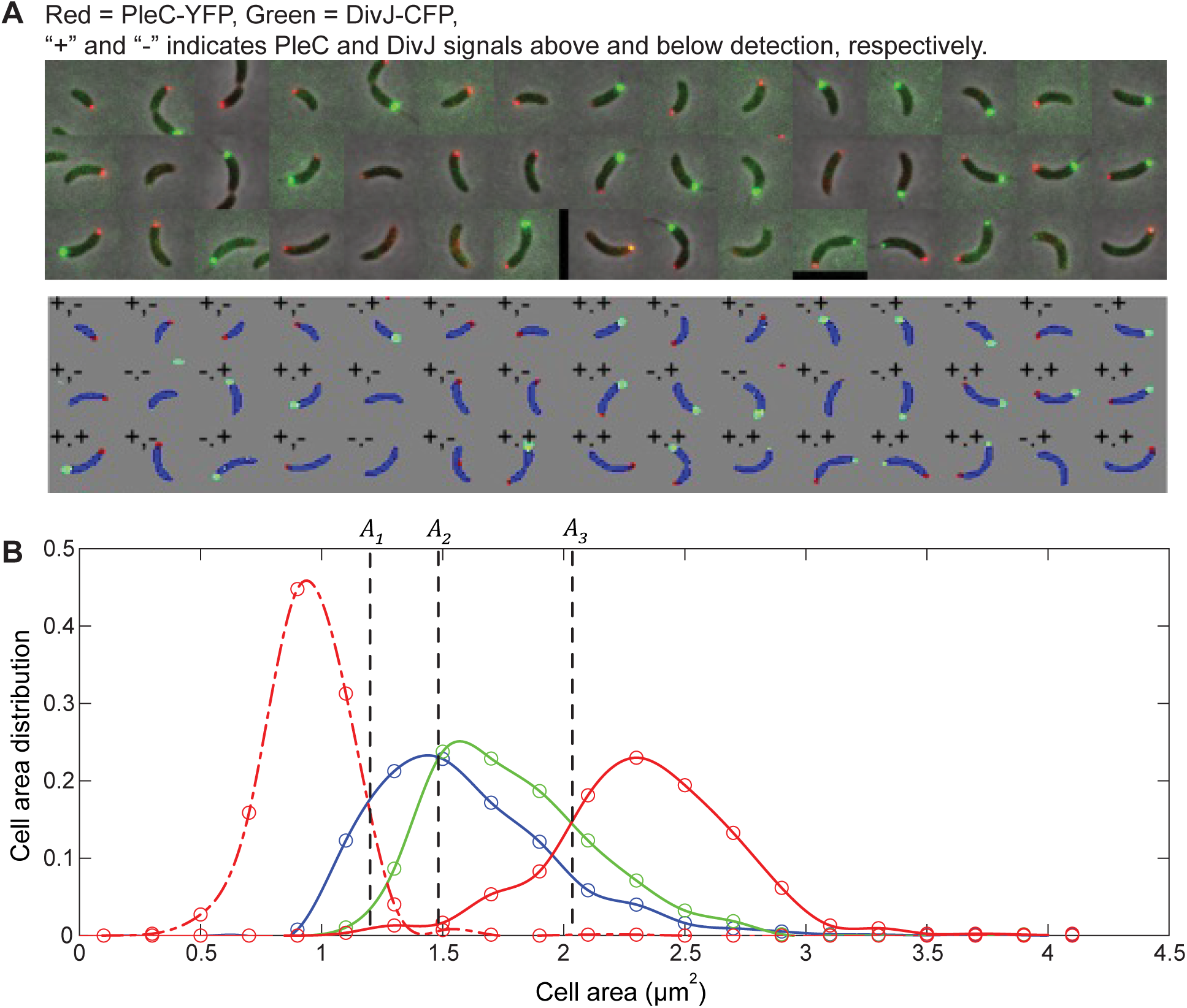
Convergent Constant Growth (CCG) model parameter extraction. (**A**) Visualization of the image analysis pipeline for the CCG model. Upper panel: Representative microscopy images of strain (LS3205) with phase and fluorescence channels merged. Phase contrast is in grey, while PleC-YFP and DivJ-CFP are pseudo-colored red and green, respectively. Cells are sorted by increasing cell area. Lower panel: the result of cell segmentation (blue region), spot detection of PleC-YFP and DivJ-CFP markers (red and green regions, respectively), and cell classification by the presence (+) or absence (-) of PleC-YFP and DivJ-CFP markers at a cell pole. (**B**) Normalized cell area distributions of swarmer (blue, 𝑔_1_(𝑠)), stalked (green, 𝑔_2_(𝑠)), and predivisional (red, 𝑔_3_(𝑠)) cells, and 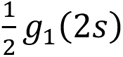 in dashed red (see Materials and Methods). The intersection of the *curves (𝐴_1_, 𝐴_2_, 𝐴_3_) provides an estimate of the average initiation and ending areas for each stage-age.

**Fig. S2.**
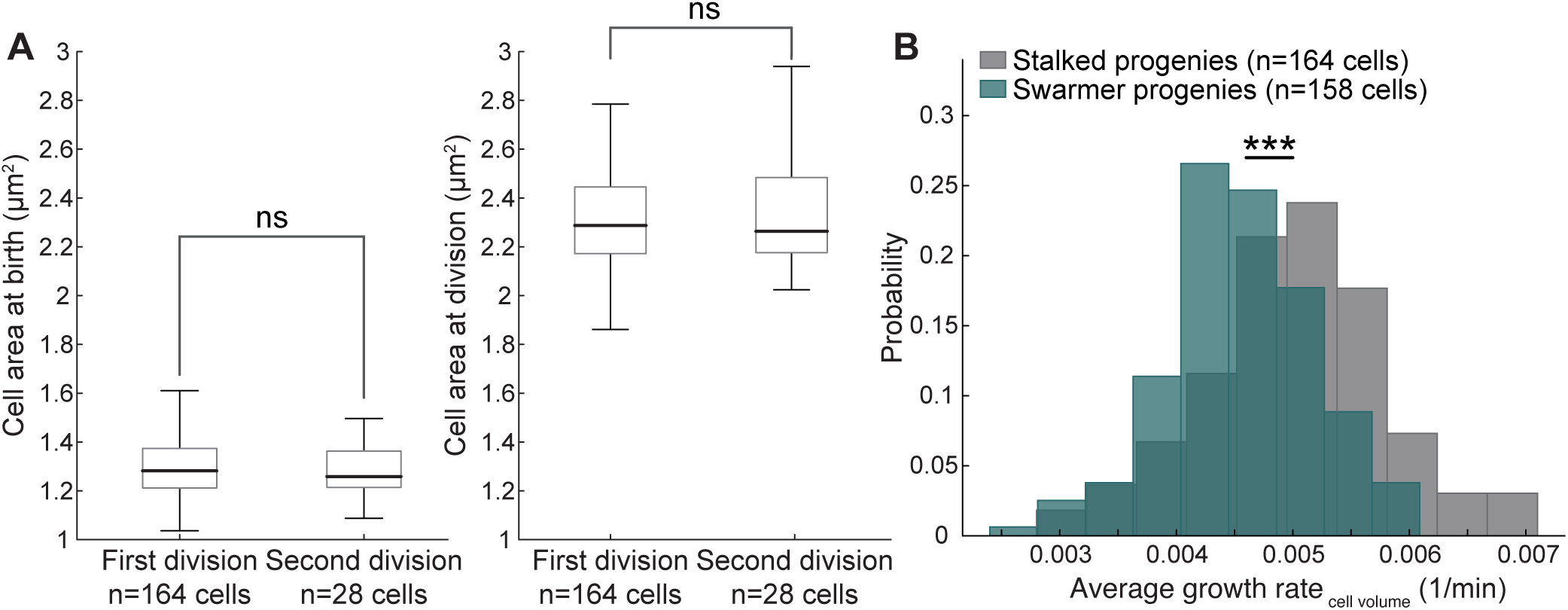
Growth analysis of *C. crescentus* from low-agarose timelapse. (**A**) Box plots showing cell area at birth (two-sample Kolmogorov-Smirnov test, p = 0.70) and division (two-sample Kolmogorov-Smirnov test, p = 0.80) for stalked progenies across two generations. Box plots show median and interquartile range for all cells sampled, whiskers show minimum and maximum observations. Cell size does not change significantly between the first and second divisions. ns indicates a statistically non-significant (p > 0.05) difference. (**B**) Distributions showing the same cell size data as Fig. 2B but with average growth rate calculated from cell volume. *** indicates p ≤ 0.001 (two-sample Kolmogorov-Smirnov test, p = 2.96 x 10^-7^).

**Fig. S3.**
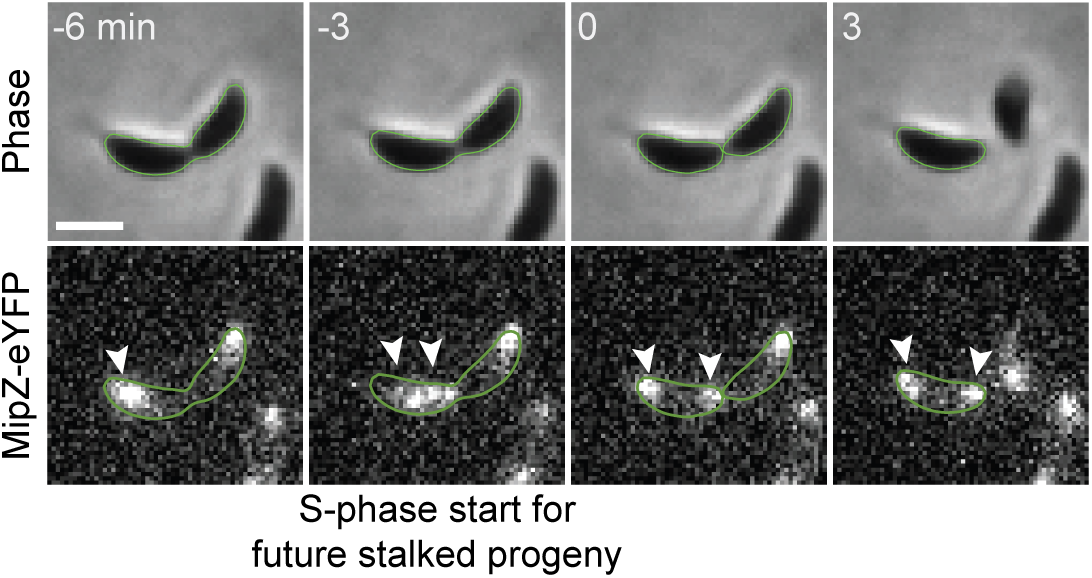
Example of DNA replication initiation in the preceding division cycle. Example montage of a stalked progeny initiating DNA replication prior to cell separation. By our definition, cell birth (time 0 min) corresponds to the frame prior to cell separation. White arrowheads show how the MipZ-eYFP spot duplicates and segregates in the future stalked progeny prior to cell separation. Scale bar represents 1 µm.

**Fig. S4.**
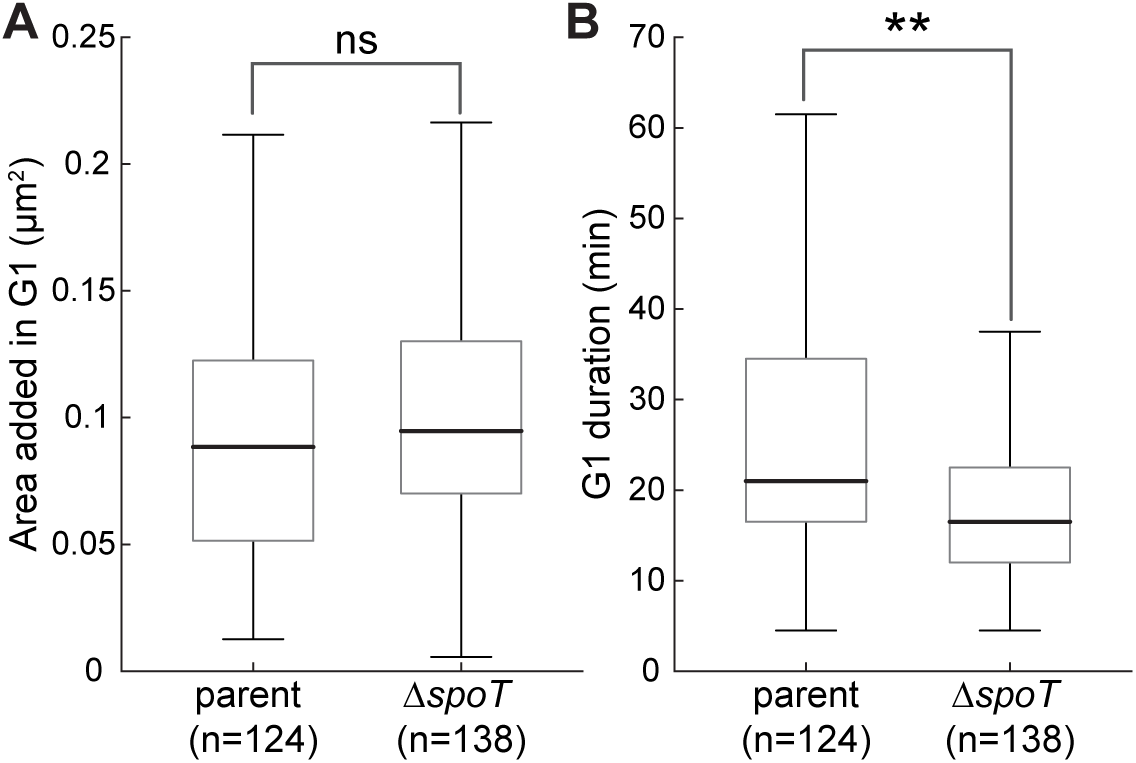
Growth rate analysis for parent and Δ*spoT* cells. Box plots show median and interquartile range for all cells sampled, whereas whiskers show minimum and maximum observations. (**A**) Box plots comparing cell area added during the G1 phase between parent (CJW2022) and Δ*spoT* (CJW7751) swarmer progenies. ns indicates a statistically non-significant (p > 0.05) difference (two-sample Kolmogorov-Smirnov test, p = 0.25). (**B**) Box plots comparing G1 durations between parent (CJW2022) and Δ*spoT* (CJW7751) swarmer progenies. ** indicates p ≤ 0.01 (two-sample Kolmogorov-Smirnov test, p = 1.9 x 10^-3^).

### Movie Legends

**Movie S1 (separate file). Representative phase contrast timelapse movie of a wildtype (CB15N) stalked and swarmer progeny growing and dividing on a low-agarose pad.** Strain is wildtype (CB15N). Scale bar represents 2 µm.

**Movie S2 (separate file). Video of low-agarose timelapse showing multiple cells growing and dividing.** Strain is wildtype (CB15N). Scale bar represents 2 µm.

**Movie S3 (separate file). Representative timelapse movie of cells bearing MipZ-eYFP and growing on low-agarose pad.** Strain is CJW2022. Left panel is phase contrast, right panel is the YFP channel. Scale bar represents 2 µm.

**Movie S4 (separate file). Example video of a stalked progeny that initiates DNA replication in its mother cell before cell separation.** Strain is CJW2022. Scale bar represents 2 µm.

